# Rapid prototyping of metabolites detection by bacterial biosensors in human fecal samples

**DOI:** 10.1101/2022.01.21.476945

**Authors:** Ana Zuñiga, Lucile Boivineau, Pauline Mayonove, Ismael Conejero, Georges-Philippe Pageaux, Romain Altwegg, Jerome Bonnet

## Abstract

Gut metabolites are pivotal mediators of host-microbiome interactions and provide an important window on human physiology and disease. However, current methods to monitor gut metabolites rely on heavy and expensive technologies such as liquid chromatography-mass spectrometry (LCMS). In that context, robust, fast, field-deployable, and cost-effective strategies for monitoring fecal metabolites would support large-scale functional studies and routine monitoring of metabolites biomarkers associated with pathological conditions. Living cells are an attractive option to engineer biosensors due to their ability to detect and process many environmental signals and their self-replicating nature. Here we optimized a protocol for feces processing and gut metabolites detection using bacterial biosensors (bactosensors), enabling rapid evaluation of their operational capacity in these samples. We show that a simple filtration step is enough to remove host microbes and reproducibly obtain a physiological-derived media retaining important characteristics of human feces, such as matrix effects and endogenous metabolites. We measured how fecal samples affect the performance of biosensors for benzoate, lactate, anhydrotetracycline, and bile acids, and found that bactosensors are highly sensitive to fecal matrices. Sensitivity to the matrix is biosensor-dependent but also varies between individuals, highlighting the need for case-by-case optimization for bactosensors operation in feces. Finally, by detecting endogenous bile acids, we demonstrate that bactosensor can be used for metabolites monitoring in feces. This work lays the foundation for the optimization and use of bacterial biosensors for fecal metabolites monitoring. In the future, our method could also allow rapid pre-prototyping of engineered bacteria designed to operate in the gut, with applications to *in situ* diagnostics and therapeutics.

## INTRODUCTION

The human gut microbiota contains a large number of interacting species of bacteria, archaea, bacteriophages, eukaryotic virus, and fungi which together create a complex ecosystem able to influence human physiology, pathologies, and behavior (Oliphant et al., 2019; Fan and Pedersen, 2020). The microbiome plays important roles in human homeostasis mostly in link with metabolism (Smith et al., 2013). Multiple studies have linked abnormal-gut microbiota with altered metabolite profiles of patients with different diseases as metabolic liver disease (Jiang et al., 2015; Schwenger et al., 2019), inflammatory bowel disease (Duboc et al., 2013; Nikolaus et al., 2017; Franzosa et al., 2019) as well as metabolic disorders like obesity and malnutrition (Ridaura et al., 2013; Smith et al., 2013; Sharon et al., 2014).

The analysis of feces metabolites has opened a new window on the complex interactions occurring within the gut (Wang et al., 2011; Patterson et al., 2016; Woting and Blaut, 2016; Jia et al., 2018). In the clinics, simple and quantitative tests enable measurements of fatty acid content or malabsorption of carbohydrates by analyzing the pH of feces (Caballero et al., 1983; Eherer and Fordtran, 1992). The detection of fecal proteins, particularly calprotectin, help diagnose and monitor inflammatory bowel diseases (Manceau et al., 2017). In addition, bacterial infections of the gut can be detected using culture-based or molecular genotyping strategies (Karu et al., 2018). Finally, mass spectrometry coupled with liquid chromatography (LC-MS) has been successfully applied to measure the levels of metabolites in human feces allowing the prediction of key association between diet and microbiome (Bjerrum et al., 2015; Bar et al., 2020). While LC-MS is increasingly used in clinical diagnosis and allows for a general and precise metabolic profiling, it is still impractical and expensive for daily monitoring of metabolites (Seger and Salzmann, 2020). Additionally, heavy methods such as LC-MS are not deployable in the field or at home, restricting large-scale prospective routine monitoring of patients. The development of innovative point-of-care (POC) testing could enable real time clinical decision-making, eliminating requirements for specialists to perform and analyze the test. Therefore, new technologies are needed to support fast, field-deployable, and cost-effective detection of the metabolites produced by the microbiome in human samples such as feces.

Programmable bacteria present an attractive technology for engineering portable biosensor devices that could help address these challenges. A biosensor is composed of a biological sensing component, which recognizes a chemical or physical change, coupled to a transducing element that produces a measurable signal in response to the environmental change (Daunert et al., 2000). A wide number of bacterial biosensors have been engineered to detect different types of analytes by connecting natural transcriptional responses to different reporter genes (Chang et al., 2017; Hicks et al., 2020). Bacterial biosensors have significant potential in applications for medical diagnosis as they perform analyte detection with a robust response, high sensitivity, and can be optimized to detect molecules in complex media such as clinical human samples (Courbet et al., 2015; Watstein and Styczynski, 2017; Chang et al., 2021). Moreover, they are inexpensive and easy to manipulate and store. Synthetic biology has enabled bacterial biosensor improvement by providing a large number of standardized genetic parts, together with systematic strategies for organism engineering, resulting in bactosensors with a higher specificity, able to detect molecules in a relevant range of concentration (Hicks et al., 2020).

In addition, synthetic biology has demonstrated the promising *in vivo* application of engineered bacteria for the treatment of disease, including metabolic disorders (Nelson et al., 2021), infections (Daeffler et al., 2017; Riglar and Silver, 2018) and modulation of tumor microenvironment (Canale et al 2021). Characterizing and optimizing the sensing performance of these bacteria under physiological conditions could improve their *in vivo* applications and their therapeutics abilities.

Here we developed a rapid prototyping method for metabolite detection in feces by bacterial biosensors. By characterizing the sensing performance of different biosensors in feces, we have assessed feces matrix effect on their response, which varies depending on the type of biosensor and the target molecule. Using this method, we were able to detect exogenous and endogenous molecules, demonstrating that the detection of fecal metabolites by bactosensors is a feasible monitoring strategy. In addition, this method could be used for rapid pre-prototyping of engineered bacteria designed to operate in the gut, with applications to *in vivo* diagnosis and therapeutics.

## MATERIAL AND METHODS

### Strains

All bactosensors strains used in this study are provided in Supporting Information (**Supplementary Table S1**). All experiments were performed using the *E. coli* strains DH5αZ1 and NEB10β (New England Biolabs). The different biosensors were grown in LB media with corresponding antibiotics (kanamycin 25 μg/mL or chloramphenicol 25 μg/mL). The inducers were: anhydrotetracycline used at a final concentration of 200 nM, benzoic acid used at a final concentration of 100 μM, L-lactate used at final concentration of 10 mM and taurocholic acid (TCA) and glycodeoxycholic acid (GDCA) used at different final concentrations. All chemicals used in this research were purchased from Sigma-Aldrich.

### Human feces samples collection

Feces samples from routine monitoring of IBD patients were obtained from the Hepatogastroenterology and Bacteriology service at CHU Montpellier (France), in accordance with ethics committee approval (# 202101009). About 100-120 mg of samples were collected by using a swab (ESWABR1, Copan ITALIA S.p.A) with 1mL of eSwab™ buffer (consisting of modified liquid Amies medium (Amies, 1967) containing sodium chloride, potassium chloride, magnesium chloride, calcium chloride, monopotassium phosphate, disodium phosphate, and sodium thioglycollate. We chose eSwab™ media as it is widely used in clinical sample collection, and has shown to help conserve both microorganisms and metabolites (Perry, 1997; Gumede et al., 2017; Saliba et al., 2020). In particular, sodium thioglycollate, which is used to maintain the reducing condition of the media, avoids oxidations of metabolites, thus contributing to the stability of metabolites and also to the growth of the cells. After resuspension in the buffer, samples were immediately stored at −80°C until use. All experiments involving feces were performed in a containment level 2 laboratory.

### Feces processing

Collected samples were defrosted and homogenized for 2 minutes by vortexing, then centrifuged at 4000 rpm for 10 min in eppendorf tubes. The supernatant was recovered in a new eppendorf tube and stored at −20°C until use. Further processing by filtering was done by using a 13 mm diameter sterile syringe filter with a 0.45 μm or 0.2 μm pore size hydrophilic PVDF membrane (Millex-HV Syringe Filter, Millipore). All feces homogenizations were done using the eSwab™ buffer. The dilution of pre-treated feces was done following the general mixing of volumes; 75 μL 2X LB medium, plus 1.5 μL of biosensor culture, plus 3 μL of inducer adjusted at the needed concentration and 70.5 μL of the feces samples diluted in eSwab™ buffer to have a final volume of 150 μL, as follow; for 10% final feces concentration: 15 μL of feces samples plus 55.5 μL of eSwab™ buffer, for 25% final feces concentration: 37.5 μL of feces samples plus 33 μL of eSwab™ buffer and for 50% dilution 75 μL of feces samples.

### Functional characterization of whole-cell biosensors in feces

Biosensor strains from glycerol stock were plated on LB agar plates supplemented with antibiotics and incubated at 37°C overnight. For functional characterization three fresh colonies of each biosensor were picked and inoculated into 0.5 mL of LB with corresponding antibiotics and grown at 37°C for 16 h in 96 DeepWell polystyrene plates (Thermo Fisher Scientific, 278606) sealed with AeraSeal film (Sigma-Aldrich, A9224-50EA) with shaking (200 rpm) and 80% of humidity in a Kuhner LT-X (Lab-Therm) incubator shaker. In the next day, the cultures were diluted 1:100 into a final volume of 150 μL of LB supplemented or not with different dilutions of feces samples, and corresponding inducers, in 96-well plates, incubated at 37°C without shaking for 16 h and analyzed by flow cytometry. Pre-treated feces samples were diluted with various volumes of eSwab homogenization buffer depending on the final target feces concentration. The cultures were incubated at 37°C for 16h without shaking, with the goal of having the simplest protocol possible for future field application to metabolite detection. Next, cells were well mixed and 100-times diluted in 1X Attune Focusing Fluid (Thermo Fisher Scientific) before cytometry analysis (for more details on the method see Figure S2).

### Enzymatic assays for total bile salts and lactate quantification in feces

Total bile acids in feces was measured using Bile Acid Assay Kit (Sigma-Aldrich MAK309, Merck, France). The L-lactate concentration was measured using a L-lactate Assay Kit (Sigma-Aldrich MAK329, Merck, France). 20 μL of pre-treated and 0.4 μm filtered feces samples were used for each reaction. All measurements were performed in duplicate in two different days.

### Flow cytometry analysis

Flow cytometry was performed on Attune NxT flow cytometer (Thermo Fisher) equipped with an autosampler and Attune NxTTM Version 2.7 Software. Experiments on Attune NxT were performed in 96-well plates with setting; FSC: 200 V, SSC: 380 V, green intensity BL1 488 nm laser and a 510/10 nm filter. All events were collected with a cutoff of 20.000 events. A control cell-line of *E. coli* containing a reference construct was grown in parallel for each experiment. This in vivo reference construct has a constitutive promoter J23101 and RBS_B0032 controlling the expression of a superfolder GFP as a reporter gene in the plasmid pSB4K5. The cells were gated based on forward and side scatter graphs and events on single-cell gates were selected and analyzed, to remove debris from the analysis, by using Flow-Jo (Treestar, Inc) software. The gating strategy is depicted in **Supplementary Figure S4**.

#### Data analysis

The calculation of relative promoter units (RPUs, (Kelly et al., 2009)) was done by normalizing the fluorescence intensity measurements of each biosensor according to the fluorescence intensity of the control cell-line *E. coli* harboring a reference construct. We quantify the geometric mean of fluorescence intensity (MFI) of the flow cytometry data and calculated RPUs according to the following equation:

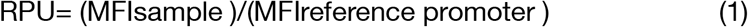

The goodness of fit and the EC50 for each data from bile acids biosensors set were calculated by applying non-linear regression using Agonist vs. response-Variable slope function using GraphPad Prism.

The relative percentage of activity of the biosensors was calculated by summing the differences of the fluorescence in RPU between the conditions with the different % of feces samples and the control condition without the feces samples, in presence of the inducer.

### RESULTS

#### Pre-treatment of human feces

To evaluate whole-cell biosensors functionality in feces, we first pre-treated samples of human feces by testing different protocols and assessing the activity of a benzoate biosensor (**Figure 1A**). Samples were collected at the hospital and immediately stored at −80°C until they were processed, as recommended to avoid changes in the metabolites and bacterial composition (Karu et al., 2018; Guan et al., 2021). Samples collection was performed in Amies medium (Amies, 1967), a derivative of Stuart’s medium (Stuart, 1956), widely used for clinical samples preservation during transportation, both in the hospital and in field experiments, making our protocol compatible with existing workflows (Perry, 1997; Gumede et al., 2017; Saliba et al., 2020). Amies medium has shown superior performance in comparative studies for microorganisms conservation and its reducing capacity helps avoid metabolite oxidations. In the context of bacterial biosensors, the compatibility of the buffer with cellular viability and growth is of particular interest. Upon measurements, the samples were defrosted, homogenized by vortexing, and centrifuged (see methods). The supernatant from 4 patients was recovered, pooled, and four pre-treatment methods were applied: (i) no filtration, meaning no change in the original microbial composition of the feces, (ii) no filtration + antibiotics, to inhibit the growth of the endogenous bacterial microbiome during the course of the experiment, (iii) filtration at 0.45 μm and (iv) filtration at 0.2 μm, to reduce the amount of host-derived microorganisms in the samples. The final pre-treated samples were diluted in culture media for a final feces concentration of 75%, 50% and 25% vol/vol (see materials and methods). We then evaluated the output signal of the biosensor in the presence or in the absence of 100 μM benzoate (**Figure 1, Supplementary Figure S1**). After 16h of induction, considerable inhibitory effects were observed in the benzoate biosensor output signal in 75% and 50% feces. On the other hand, 25% pre-treated feces samples allowed a better sensing performance. In addition, a higher output signal was observed in samples filtered with 0.45 μm and 0.2 μm compared with the non-filtered ones (**Figure 1B**). A fraction of cells with no detectable fluorescence was observable in non-filtered feces, probably corresponding to endogenous host microorganisms present in the matrix (**Figure 1B and Supplementary Figure S1**). Since the filtering of samples at 0.2 μm did not show a significant improvement from the filtered at 0.4 μm, we chose the filtering at 0.4 μm as a final step of the pre-treatment protocol of feces. These results demonstrate that feces samples have significant inhibitory matrix effects on bacterial biosensors. Filtering and diluting fecal samples allow bacterial biosensors to operate and detect metabolites in the presence of feces.

**Figure 1.**
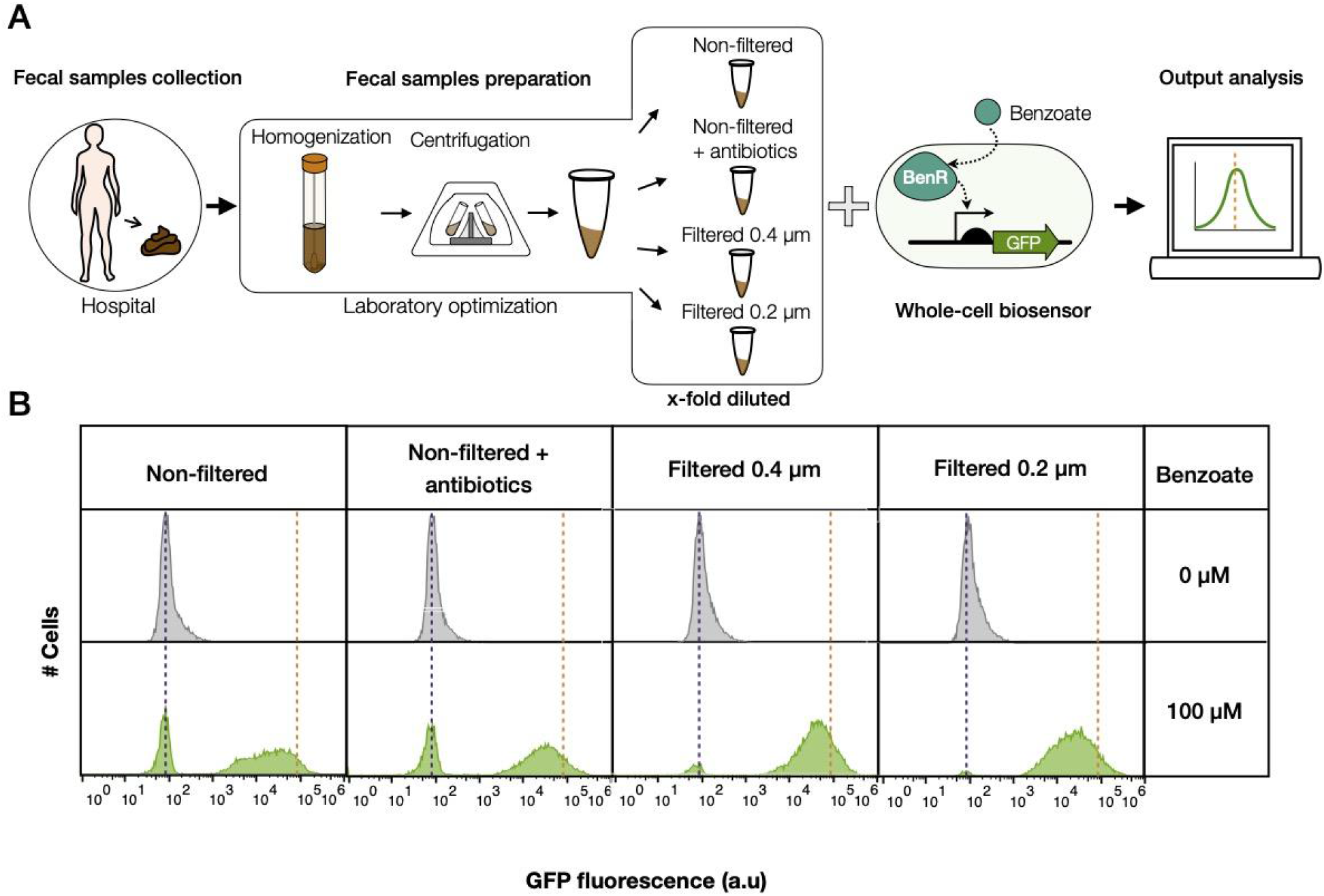
Optimization of human feces pre-treatment to reduce matrix effects. (**A**) Schematic workflow of the fecal matter treatment to assess metabolites detection by bacterial biosensors. (**B**) Samples were defrosted, homogenized and centrifuged before evaluating the pBEN biosensor performance on different pre-treated feces samples. Feces were diluted 4-fold. Cells containing the pBEN biosensor were induced or not with 100 μM benzoate and incubated at 37 °C without shaking for 16h. Dotted lines represent the mean fluorescence produced by the biosensor growing in LB only, purple without benzoate, orange with 100 μM of benzoate. Each histogram is representative of two independent experiments measured by flow cytometry.

**Figure 2.**
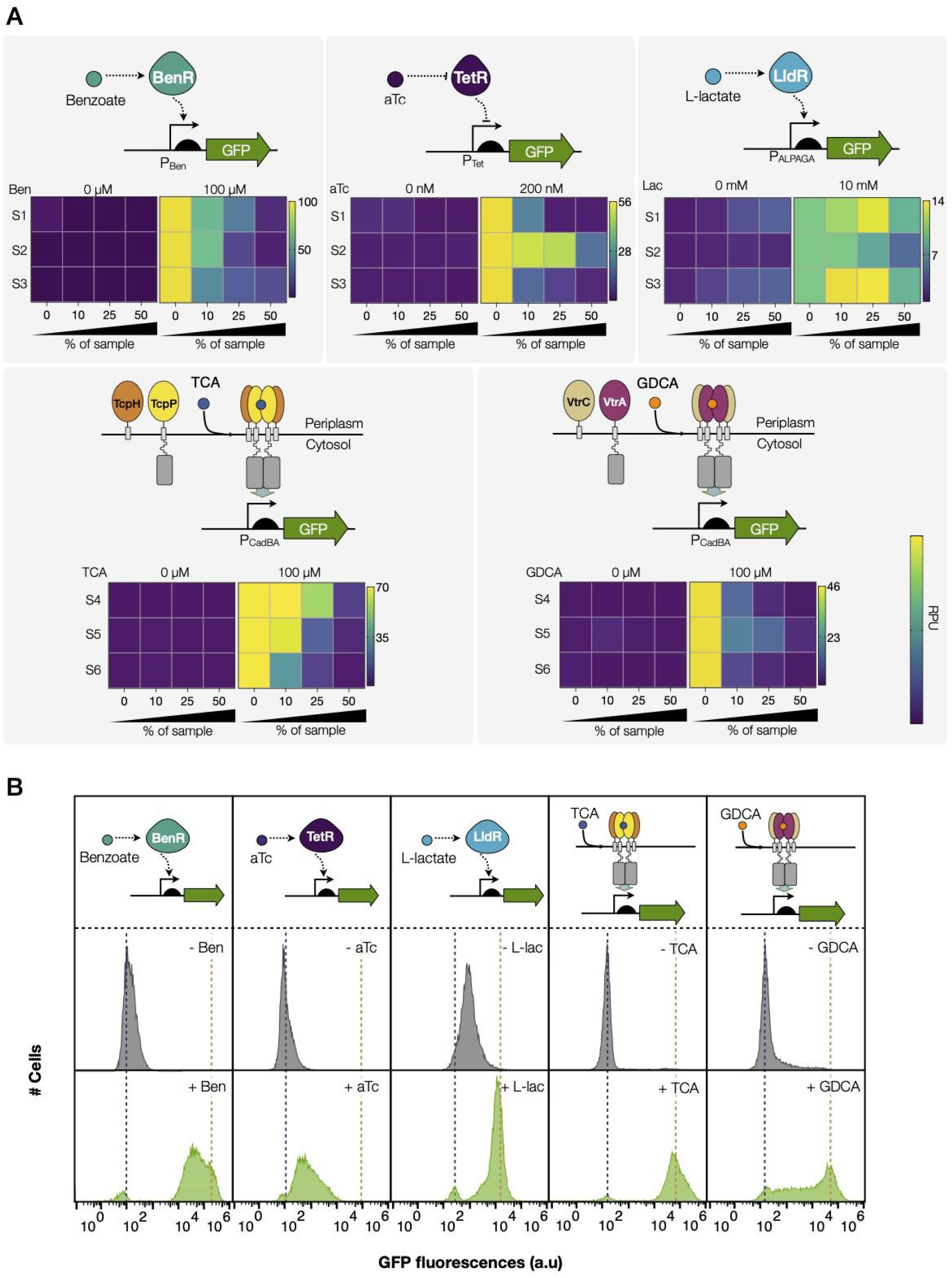
Performance of different bacterial biosensors in human feces. The matrix effects of pre-treated human feces on the performance of five different biosensors were evaluated. Three of them (*top*) are based on cytosolic transcription factor systems: the BenR activator and the pBEN promoter; the TetR repressor and the pTET promoter and the LldR regulator and the pALPAGA promoter, induced with 100 μM benzoate, 200 nM aTc and 10 mM L-lactate, respectively. The other two biosensors (*bottom*) are based on transmembrane receptors activated via ligand-induced dimerization, responding to bile acids. In the presence of 100 μM taurocholic acid (TCA), TcpP dimerizes and forms a stable complex with TcpH to activate downstream expression of the GFP reporter (*left bottom*). Glycodeoxycholic acid (GDCA, 100 μM) binds to VtrA/VtrC heterodimeric complex and activates downstream expression of the GFP reporter (*right bottom*). (**A**) Pre-treated feces samples from three different patients were used to evaluate the effect matrix on each biosensor performance. Three different dilutions of samples were used at final percentages; 10%, 25%, and 50% feces. Fluorescence intensities are expressed in relative promoter units (RPU) (see methods). The mean of three independent experiments performed in duplicate is plotted. The averages and standard deviations for these data are available in Table 1 and in supplementary data. For facilitating readability, note that the color scale was adjusted individually for each sensor with the maximum value corresponding to values measured without the addition of fecal samples. (**B**) Representative histogram showing the fluorescence of the reporter gene for each biosensor expressed as a result of different induction conditions in 25% feces and measured by flow cytometry. Inducers; 100 μM benzoate, 200 nM aTc, 10 mM L-lactate, 100 μM TCA and 100 μM GDCA.

#### Performance of bacterial biosensors in human feces

We then evaluated the sensing performance of different whole-cell biosensors in presence of pre-treated feces. We tested five different bacterial biosensors for exogenous and endogenous molecules. Three of the biosensors are based on cytosolic transcription factor systems: a biosensor responding to benzoate, based on BenR activator and the pBEN promoter (Libis et al., 2016; Zúñiga et al., 2020); a biosensor responding to aTc, based on TetR repressor and the pTET promote (Lutz and Bujard, 1997; Courbet et al., 2015), and a biosensor responding to L-lactate, based on LldR regulator and the pALPAGA promoter (Zúñiga et al., 2021). The other two biosensors, TcpPH and VtrAC, are based on synthetic transmembrane bile salts receptors activated via ligand induced dimerization (Chang et al., 2021). TcpP dimerizes and forms a stable complex with TcpH to activate downstream expression of the GFP reporter in response principally to primary bile acids, taurocholic acid (TCA) and glycochenodeoxycholic acid (GCDCA). VtrA and VtrC forms a heterodimeric complex and trigger transcription in response principally to secondary bile acids, glycodeoxycholic acid (GDCA) and taurodeoxycholic acid (TDCA) (Chang et al., 2021). We evaluated sample matrix effects on the sensing performance of these biosensors by adding pre-treated feces samples from three different individual patients at 2, 4 and 10-fold dilutions (50%, 20%, and 10% feces respectively, **Figure 3A**, see materials and methods and **Supplementary Figure S2**). For TcpP/H and VtrA/C, in order to avoid interference from endogenous cognate ligands on the biosensor, the samples were chosen for their low content in total bile acids, as measured by an enzymatic assay (**Supplementary Table S2**). All biosensors were able to detect their specific inducers in 10% feces samples. However, most of them failed to produce fluorescence in 50% feces, except for lactate biosensors that showed a higher fluorescence when higher concentration of feces was added, due to the endogenous presence of lactate in the samples (**Supplementary Table S2**). Indeed, a feces sample with low measured lactate concentration (**Figure 3A**, patient S3) did not exhibit such increase in fluorescence. The different sensors exhibited diverse degrees of sensitivity to fecal matrices (**Figure 3B, Supplementary Figure S3, Table 1 and Supplementary Table S3**). TetR was strongly inhibited (more than 50%), even in 10% feces. Interestingly, the VtrAC system exhibited a non-homogeneous response and was strongly inhibited too, although relying on a similar architecture than TcpPH, which was not significantly affected. This difference might be due to the fact that the behavior of TcpPH was previously optimized by circuit tuning and directed evolution (Chang et al., 2021).

**Table 1.** Functional analysis of biosensors in human feces

| % Of Sample | 0 % |  |  | 10 % |  |  | 25 % |  |  | 50 % |  |  |
| --- | --- | --- | --- | --- | --- | --- | --- | --- | --- | --- | --- | --- |
| Inducer | Leakage RPU | Max Fold Change | Max Swing RPU | Leakage RPU | Max Fold Change | Max Swing RPU | Leakage RPU | Max Fold Change | Max Swing RPU | Leakage RPU | Max Fold Change | Max Swing RPU |
| Benzoate | 0.22 ± 0 | 104 ± 25 | 103 ± 14 | 0.21 ± 0 | 56 ± 13 | 56 ± 8 | 0.22 ± 0 | 23 ± 10 | 23 ± 5 | 0.2 ± 0 | 8.5 ± 6 | 8.2 ± 3 |
| Anhydrotetracycline (aTc) | 2.3 ± 1.3 | 27 ± 11 | 55 ± 10 | 2.1 ± 1.7 | 22 ± 10 | 42 ± 16 | 0.9 ± 0.6 | 14 ± 12 | 18 ± 10 | 0.8 ± 0.7 | 8 ± 5 | 6 ± 4 |
| L-lactate | 1.3 ± 0 | 8 ± 0.2 | 8.7 ± 0 | 2 ± 0.6 | 6 ± 1.2 | 10 ± 1 | 3 ± 0.4 | 4 ± 0.6 | 9 ± 1.8 | 3.4 ± 0.2 | 2 ± 0.4 | 4 ± 1.4 |
| Taurocholic acid (TCA) | 0.6 ± 0.2 | 100 ± 23 | 68 ± 4 | 1.1 ± 0.6 | 48 ± 19 | 54 ± 12 | 0.7 ± 0.3 | 36 ± 21 | 26 ± 14 | 0.6 ± 0.2 | 8 ± 3 | 4.4 ± 2 |
| Glycodeoxycholic acid (GDCA) | 1.1 ± 0.5 | 43 ± 14 | 46 ± 8 | 1.4 ± 1.8 | 8.8 ± 6 | 11 ± 2 | 0.6 ± 0.4 | 10 ± 7 | 6 ± 3 | 0.5 ± 0.2 | 4.3 ± 2 | 1.6 ± 0.8 |

RPU: reference promoter units. The leakage RPU, measured in a non-induced state. The Max Fold change corresponds to the fold change between the induced state and the non-induced state. The Max Swing RPU corresponds to the subtraction of RPU between the induced state and the non-induced state. The average of three independent experiments and standard deviations ( ± ) are indicated. The concentration of inducers were; 100 μM Benzoate, 200 nM aTc, 10mM L-lactate, 100 μM TCA, 100 μM GDCA.

**Figure 3.**
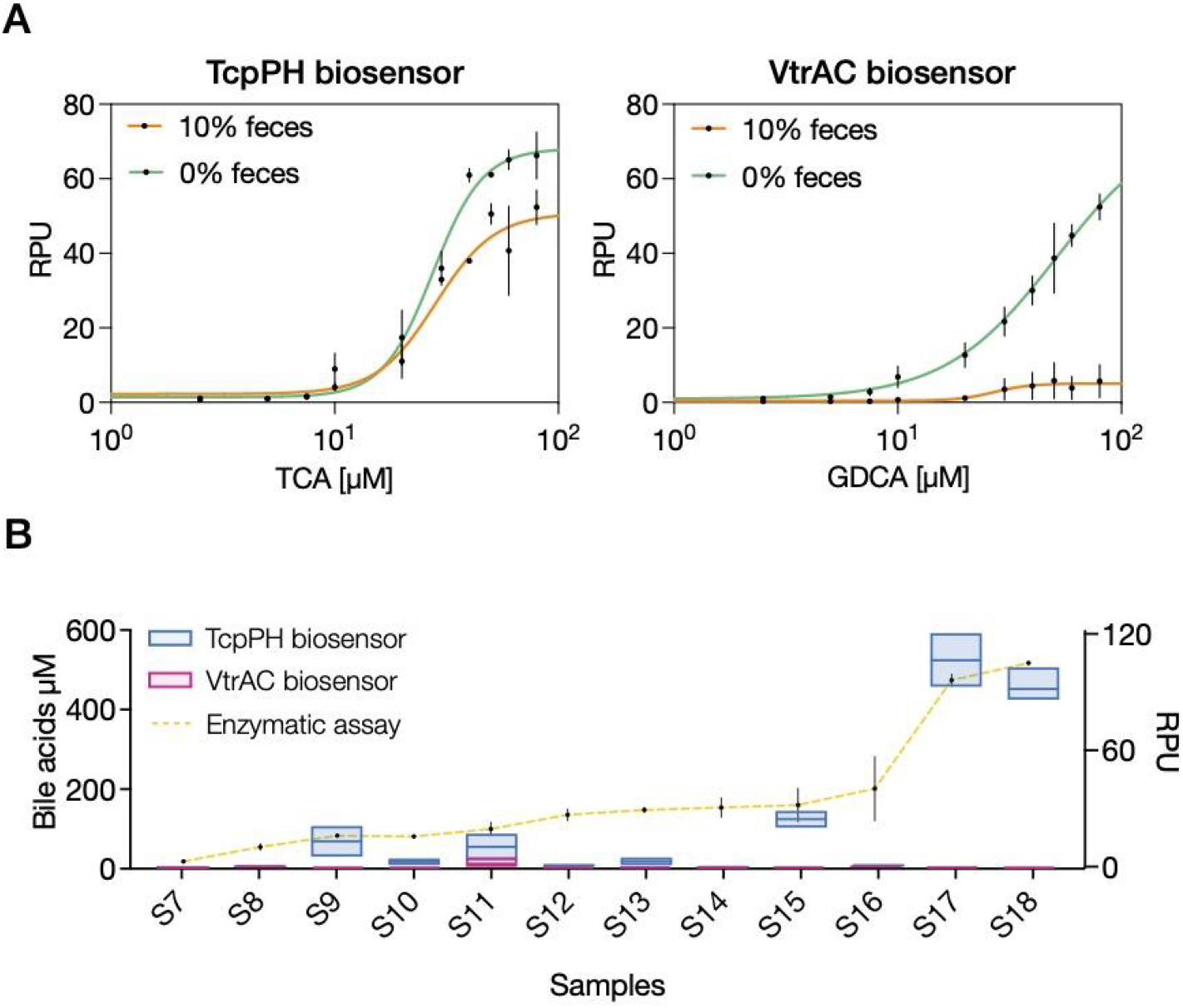
Bile acids detection in pre-treated feces samples. (**A**) Response function of TcpPH biosensor (left) and VtrAC biosensor (right) to spiked taurocholic acid (TCA) and glycodeoxycholic acid (GDCA) in the presence of 10-fold diluted feces (Three different samples were pooled; samples S5, S6 and S7). Data points correspond to the mean value of four replicates on four different days. Error bars: ±SD. (**B**) Comparison of total bile acid detection between TcpPH (blue square) and VtrAC (red square) biosensors, right axis, and enzymatic assay (yellow line) left axis. Samples were ordered according to their total bile salts concentration measured by the enzymatic assay. Data points correspond to the mean value of three replicates performed in duplicate on three different days. Error bars: ±SD.

**Figure 4.**
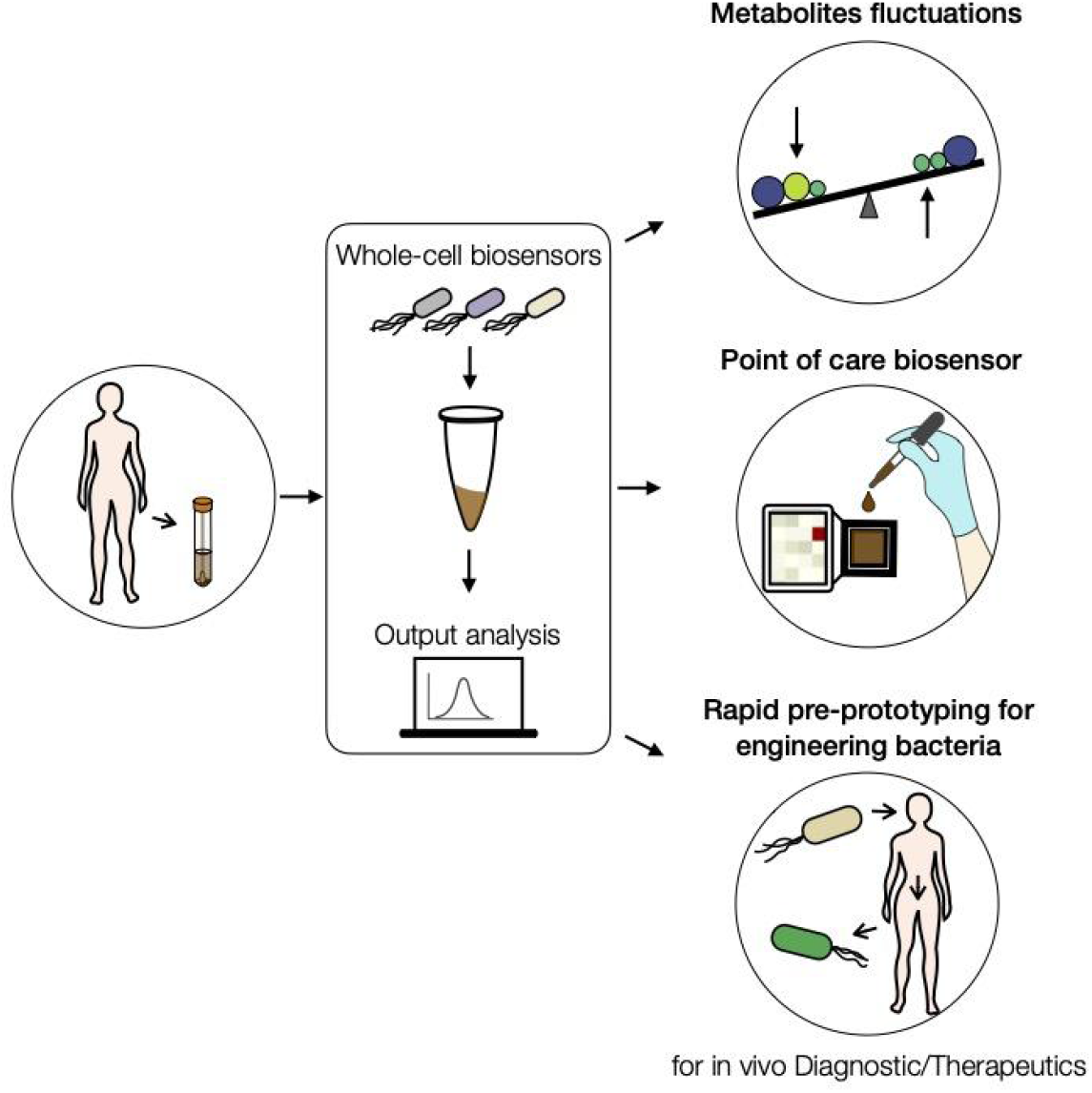
Summary of future applications for rapid prototyping of bactosensors and metabolites detection in human fecal samples. By applying this method, human feces could be collected and homogenized at the hospital, and directly used to measure the levels of targeted metabolites by using bacterial biosensors. These measurements will help to understand metabolite fluctuations under a particular diet intake, for example. Similarly, this strategy would allow the engineering of point-of-care biosensors for performing medical diagnostics by measuring a set of targeted metabolites in feces. Finally, this prototyping approach could also support the engineering of therapeutic bacteria by allowing a fast characterization and optimization of their sensing performance closer to physiological conditions in fecal matrices.

Importantly, a clear patient-to-patient variability was observed for every biosensor, suggesting that feces matrix effects responsible for biosensor inhibition could be due to specific factors having varying abundance in different patients. **In all, these results show that several bacterial biosensors can operate in feces samples, yet have diverse sensitivity to fecal matrices, requiring case-by-case optimization**.

#### Detection of endogenous bile acids in human feces

Our next goal was to evaluate if using our protocol, whole-cell biosensors could be used to detect endogenous metabolites in patients’ fecal samples. As a demo, we chose to assess bile salts concentration using the TcpPH and the VtrAC sensors. Bile salts are key components of bile and are critical for fat absorption during digestion, lipids and cholesterol metabolism, host-microbe interaction and other regulatory pathways in humans (Staley et al., 2017). Around 95% of the bile salts are reabsorbed and recycled via enterohepatic cycling, while the other 5% goes to the colon where it can be converted from primary to secondary bile salts by bacterial biotransformation. Patients with inflammatory bowel diseases (IBD) have altered faecal bile salts profiles (Duboc et al., 2013; Torres et al., 2018; Lavelle and Sokol, 2020), with lower levels of secondary bile salts but higher levels of primary and 3-OH-sulfated acids (Jansson et al., 2009; Jacobs et al., 2016; Franzosa et al., 2019).

Here we aimed to measure endogenous bile acids in pre-treated samples of human feces coming from twelve different IBD patients. We first performed a dose-response curve to bile salts for TcpPH and VtrAC sensors in the presence or absence of 10% fecal samples (**Figure 3A**).

As previously, we selected samples with low bile salts concentrations to avoid interference from endogenous ligands. The TcpPH biosensor responded remarkably to spiked TCA in 10% feces, with high fold change and comparable limit of detection to the sensor operating in the absence of feces. On the other hand, and as observed previously, the VtrAC sensor was strongly inhibited by the presence of feces. TcpPH and VtrAC were then tested for endogenous bile salts detection in twelve feces samples whose total bile salts concentration had been previously measured using a commercial enzymatic assay (**Figure 3B**). Samples exhibited different bile salts concentration, with two of them (Samples S17 and S18) having very high bile salts concentrations (>500uM). As expected, the VtrAC sensor was unable to detect endogenous bile salts in any of the samples. On the other hand, the TcpP system produced a strong response in two samples, S17 and S18, having a very high bile salts concentration. For the other samples, we observed a variable response from the biosensor even if the concentrations measured via the enzymatic assay were close **(Figure 3B**, samples S8 to S16). These discrepancies might be due to the fact that the enzymatic assay quantifies total bile salts, while the TcpPH system responds more specifically to primary bile salts, such as TCA. **In conclusion, these results show that our method supports detection of endogenous metabolites by bacterial biosensors in human feces samples.**

### DISCUSSION

In this work we provide an optimized method for prototyping bacterial biosensors in human feces. We show that a simple filtration step is enough to remove host microbes and reproducibly obtain a physiological-derived media retaining important characteristics of human feces, such as matrix effects and endogenous metabolites (e.g. bile acids). We found significant inhibitory matrix effects of feces on the bactosensors tested, although the sensitivity of the different biosensors to fecal samples varied. In addition, matrix effects varied significantly from patient to patient. This patient-to-patient variability could be due to host or microbiome derived molecules that interfere with biosensor physiology. It is also possible that some medications inhibit the sensors. A detailed knowledge of patients’ full clinical picture and current treatments will be important to interpret the sensor’s response. In all, the biosensors tested here are highly sensitive to fecal samples, and the optimal working conditions in our studies were general at a 10% feces dilution.

These results highlight two important points for future bactosensor prototyping in feces: (i) because of sensor-specific sensitivity to fecal matrices, case-by-case optimization of every new sensor for operation in feces is required and (ii) testing the sensor over various individual samples coming from different patients is critical to obtain a sensor working over a wide range of real-world conditions. Thanks to our methodology, these characterizations will be simplified.

Interestingly, the TcpPH bile salts biosensor was capable of detecting high concentrations of endogenous bile salts in samples from several patients, in accordance with enzymatic measurements. As a proof of concept, these data demonstrate, for the first time to our knowledge, the possibility of using bacterial biosensors to detect endogenous metabolites in human feces.

How could bacterial biosensors operating in fecal samples be optimized in the future? First, other reporters having a higher signal to noise ratio, such as luciferase, might be evaluated. Yet, unless using the *lux*CDABE operon, which has lower performance, optimized luciferase systems such as nanoluc, while providing a lower limit-of-detection, work better after cell lysis, which would complicate the assay protocol (Lopreside et al., 2019). Second, amplifying genetic devices such as recombinase switches or hrp transcription factors might help combat matrix effects and enable operation at higher concentration, thereby supporting lower limits of detection (Courbet et al., 2015; Wan et al., 2019). Recombinase-mediated inversion or excision could also allow *post facto* analysis of biomarker presence through DNA sequencing or PCR (Courbet et al., 2015).

The method shown here could be performed on a lab-on-chip device enabling successive feces samples filtration, dilution and sensing assay in an automated manner (Wu et al., 2017, 2018; Arshavsky-Graham and Segal, 2020). Such devices would open the door to field-deployable, point-of-care gut metabolite detection either for diagnostics or epidemiological purposes.

Finally, another potential and compelling application of our method is its use for rapid and simple prototyping of engineered “smart” gut probiotics. Engineered bacteria have recently been developed to detect and/or treat many pathological conditions such as inflammation, diabetes, phenylketonuria, hyperammonemia, and cancer (Riglar and Silver, 2018). As of now, these strains have been evaluated in animal or in cellular co-culture models (Mimee et al., 2016; Daeffler et al., 2017; Taketani et al., 2020; Nelson et al., 2021). While providing valuable information, these models present limitations in terms of time, physiological relevance, and amenability to screening. The use of human fecal samples could complement these approaches by providing a fast and efficient method to assess the matrix effects of fecal matter on bacterial sensors and therapeutics and optimize their behavior.

## Supporting information

Raw data

DNA sequences for constructs

## ACKNOWLEDGMENTS

We thank members of the synthetic biology group and of the CBS for fruitful discussions and feedback. We are grateful to the patients for participating in this study and providing their samples, and to the personnel of the Montpellier CHU hospital for collecting the samples. This work was supported by an ERC starting grant “COMPUCELL” and an ANR “SynbioDiag” to J.B. J.B. also acknowledges the INSERM Atip-Avenir program and the Bettencourt-Schueller Foundation for continuous support. The CBS acknowledges support from the French Infrastructure for Integrated Structural Biology (FRISBI) ANR-10-INSB-05-01.

## AUTHORS CONTRIBUTION

A.Z., I.C. and J.B. designed the research. A.Z. designed and performed all the experiments. P.M contributed to perform the experiments at the L2 laboratory. L.B., G.-P.P. and R.A. collected clinical samples. A.Z. and J.B. analyzed the data. A.Z. and J.B. wrote the article. All authors reviewed and approved the manuscript.

