## Supplementary figures for "Rapid prototyping of metabolites detection by bacterial biosensors in human fecal samples"

Supplementary Figures S1-S4

Supplementary Tables S1-S3

**Supplementary Figure S1**

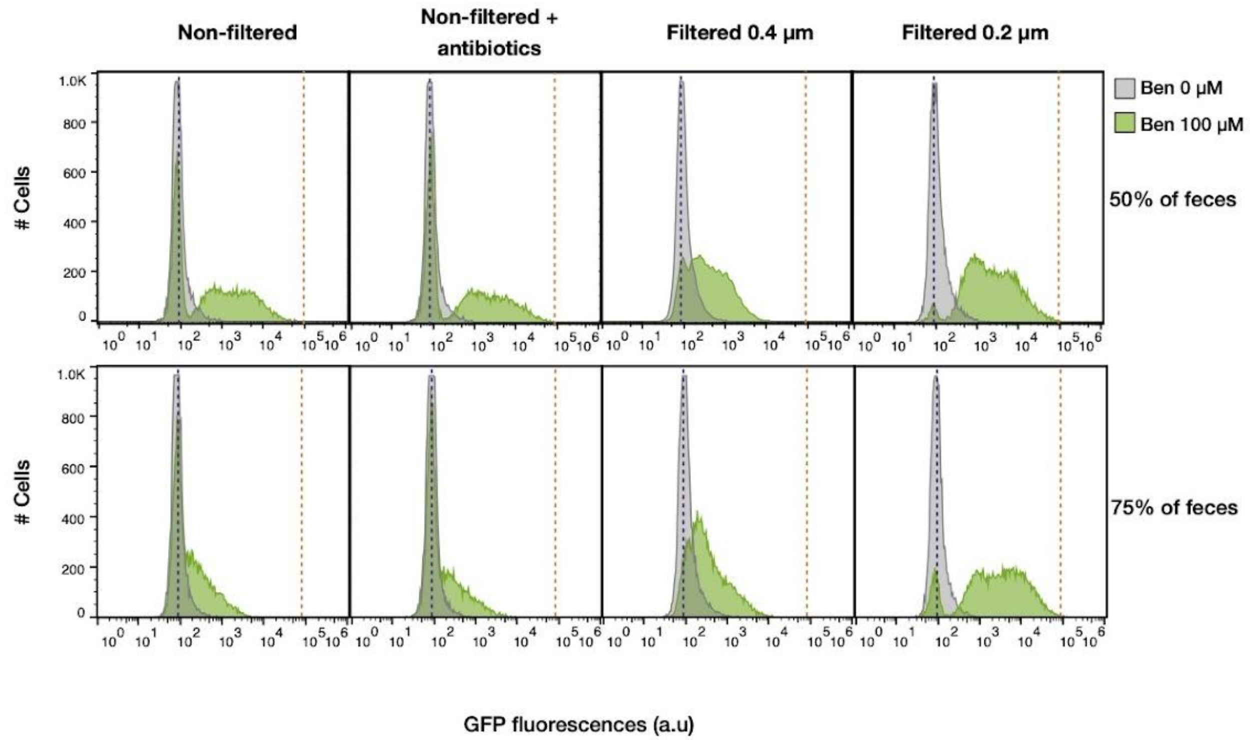


**Supplementary Figure S1. Optimization of human feces pre-treatment for a low matrix effect.** Samples were defrosted, homogenized and centrifuged before evaluating pBEN biosensor performance on different pre-treated feces. Feces were diluted 2- and 1.3-fold on LB (50% and 75%, respectively). Cells corresponding to pBEN biosensor were induced or not with benzoate 100 µM and incubated at 37 °C without shaking for 16h. Dot lines represent the mean fluorescence produced by the biosensor growing in LB only, purple without benzoate, orange with 100 µM of benzoate. Each histogram shows fluorescent reporter genes expressed as a result of different induction conditions. Each histogram is representative of two different experiments measured by flow cytometry.

**Supplementary figure S2**

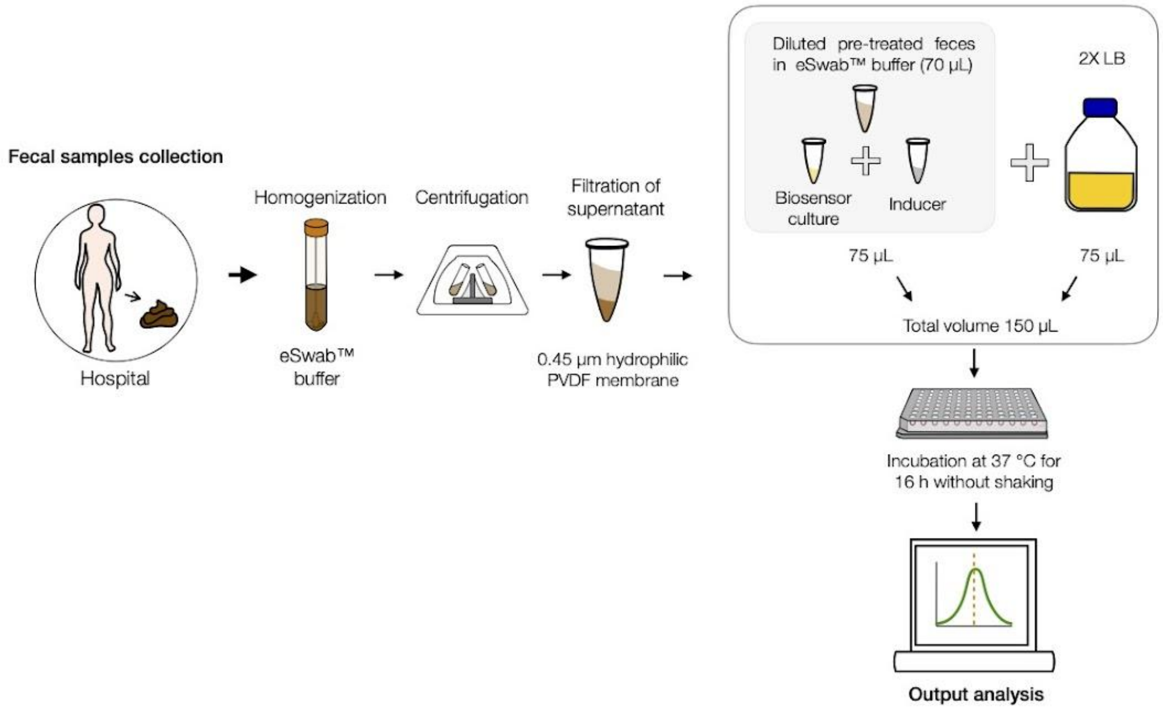


**Supplementary figure S2. Methodology for metabolites detection by bacterial biosensors in fecal samples.** Samples collected at the hospital are homogenized in the commercial eSwab buffer, then centrifuged for 10 min at 8.000 rpm, the supernatant collected and filtered by 0.45 µm hydrophilic PVDF membrane filter. Next, these pre-treated samples are mixed with 75 µL of 2X LB medium, eSwab homogenization buffer, biosensor culture and inducer until a final volume of 150 µL. The cultures are incubated at 37 °C without shaking for 16h, before cytometry analysis.

**Supplementary figure S3**

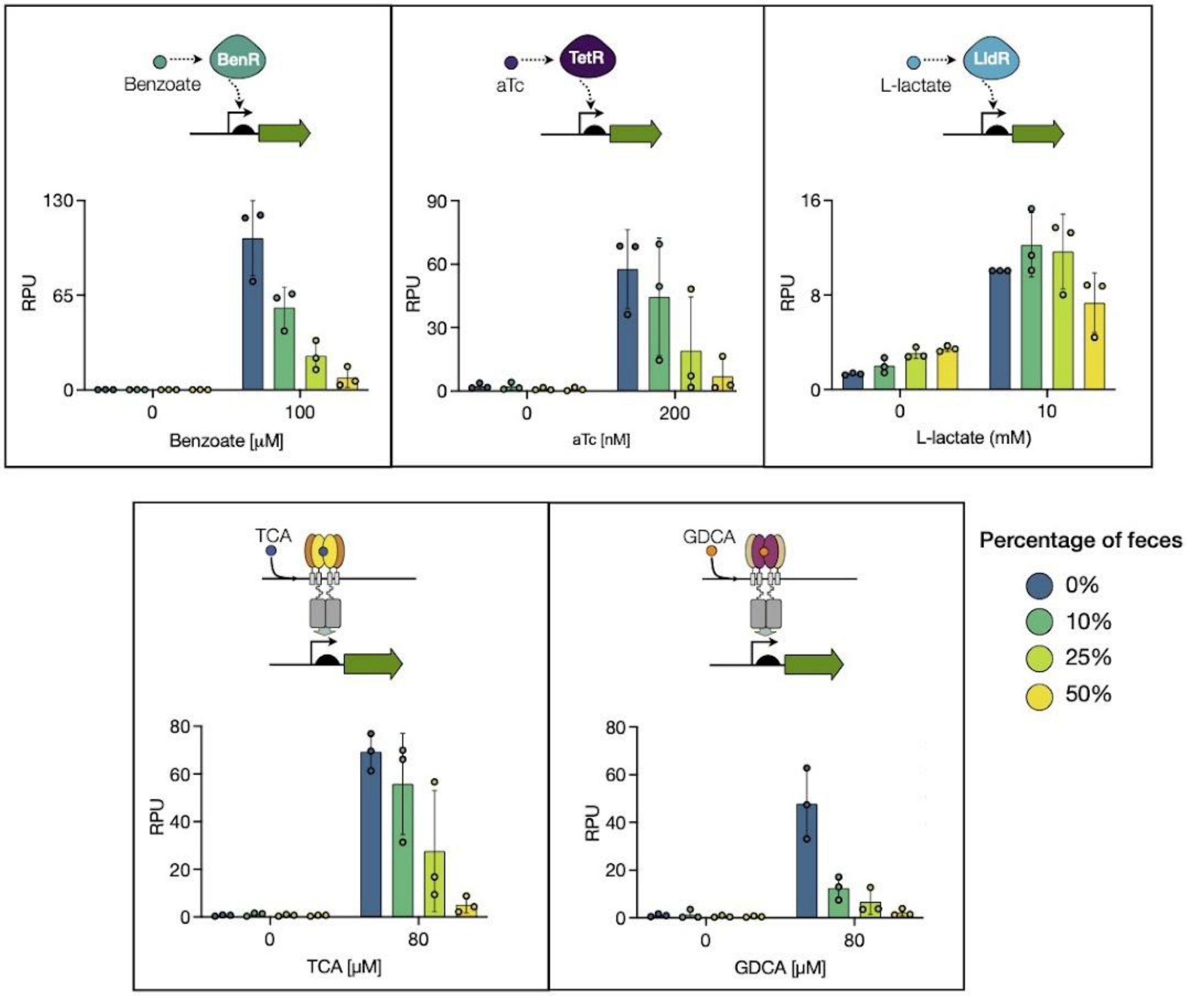


**Supplementary figure S3. Matrix effect of pre-treated human feces on the performance of five different biosensors.** Pre-treated samples of feces from three different patients were used to evaluate the effect matrix on each biosensor performance. Three different dilutions of samples were used at final percentages; 10%, 25%, 50%. The relative promoter units (RPU) for each biosensor is shown. The geometric mean of two technical replicates performed on different days is plotted. The averages and standard deviations for these data are provided in supplementary excel data. Inducers; 100 µM benzoate, 200 nM aTc, 10 mM L-lactate, 100 µM TCA and 100 µM GDCA.

**Supplementary figure S4**

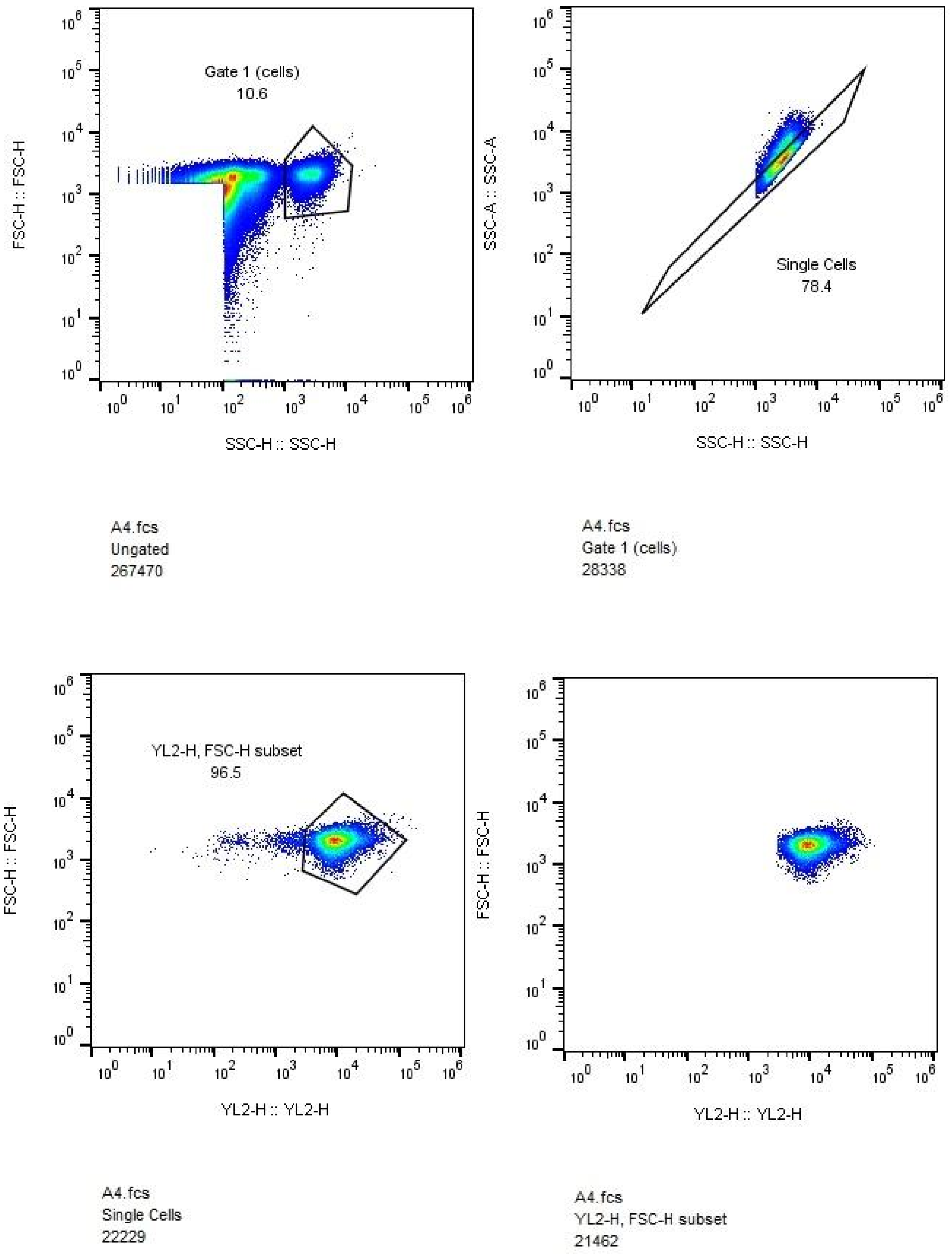


**Supplementary Figure S4.** Figure exemplifying the gating strategy. Gates were designed based on FSC-H vs SSC-H graphs to remove debris from the analysis and SSC-A vs SSC-H to doublet discrimination (lef and right upper panel). Cell subsets expressing the same fluorescent protein were selected with gates based on green, red and blue intensities (bottom panels).

Table S1. Bacterial biosensor used in this study

| Bacterial Biosensor | Strain | Plasmid | ORI | Reference |
|---|---|---|---|---|
| **TcpP/H** | *E. coli* NEB10β | J64100 $Chl^r$ | p15A | Chang et al., 2021 |
| **VtrA/C** | *E. coli* NEB10β | J64100 $Chl^r$ | p15A | Chang et al., 2021 |
| **TetR-pTET** | *E. coli* DH5αZ1 | pZ derivative $Km^r$ | p15A | Lutz and Bujard, 1997 |
| **LldR-pALPAGA** | *E. coli* DH5αZ1 | pSB4K5 $Km^r$ | pSC101 | Zúñiga et al., 2021 |
| **BenR-pBEN** | *E. coli* DH5αZ1 | J64100 $Chl^r$ | ColE1 | Zúñiga et al., 2020 |

$Chl^r$, $Km^r$ : Resistance to chloramphenicol and kanamycin, respectively.

Table S2. Bile acid and L-lactate levels in feces samples

| Sample number | Total bile acids (μM) | L-lactate (mM) |
|---|---|---|
| **S1** | 124 | 1.6 |
| **S2** | 68.6 | 0.1 |
| **S3** | 63.8 | 2.0 |
| **S4** | 29.6 | 1.1 |
| **S5** | 15.2 | 1.0 |
| **S6** | 13.5 | 0.6 |
| **S7** | 29 | |
| **S8** | 67 | |
| **S9** | 96.5 | |
| **S10** | 94 | |
| **S11** | 113.3 | |
| **S12** | 150.7 | |
| **S13** | 163.4 | |
| **S14** | 170 | |
| **S15** | 176 | |
| **S16** | 219 | |
| **S17** | 501 | |
| **S18** | 548 | |

Values were determined by enzymatic assays.

*Relative percentage of activity (RPA)= $100-((\sum(RPU_{is})-(RPU_{ic})/\sum(RPU_{ic}))*100)$ where $RPU_{is}$ correspond to the fluorescence intensity in RPU in presence of different % of feces and the inductor and $RPU_{ic}$ correspond to the fluorescence intensity in RPU in control condition in presence only of inductor.
